# The genomic basis of the plant island syndrome in Darwin’s giant daisies

**DOI:** 10.1101/2022.01.26.477903

**Authors:** José Cerca, Bent Petersen, José Miguel Lazaro Guevara, Angel Rivera-Colón, Siri Birkeland, Joel Vizueta, Siyu Li, João Loureiro, Chatchai Kosawang, Patricia Jaramillo Díaz, Gonzalo Rivas-Torres, Mario Fernández-Mazuecos, Pablo Vargas, Ross McCauley, Gitte Petersen, Luisa Santos-Bay, Nathan Wales, Julian Catchen, Daniel Machado, Michael D. Nowak, Alexander Suh, Neelima Sinha, Lene R. Nielsen, Ole Seberg, M. Thomas P. Gilbert, James H. Leebens-Mack, Loren Rieseberg, Michael D. Martin

## Abstract

Oceanic archipelagos comprise multiple disparate environments over small geographic areas and are isolated from other biotas. These conditions have led to some of the most spectacular adaptive radiations, which have been key to our understanding of evolution, and offer a unique chance to characterise the genomic basis underlying rapid and pronounced phenotypic changes. Repeated patterns of evolutionary change in plants on oceanic archipelagos, i.e. the plant island syndrome, include changes in leaf morphology, acquisition of perennial life-style, and change of ploidy. Here, we describe the genome of the critically endangered and Galápagos endemic *Scalesia atractyloides* Arnot., obtaining a chromosome-resolved 3.2-Gbp assembly with 43,093 candidate gene models. Using a combination of fossil transposable elements, *k*-mer spectra analyses and orthologue assignment, we identify the two ancestral subgenomes and date their divergence and the polyploidization event, concluding that the ancestor of all *Scalesia* species on the Galápagos was an allotetraploid. There are a comparable number of genes and transposable elements across the two subgenomes, and while their synteny has been mostly conserved, we find multiple inversions that may have facilitated adaptation. We identify clear signatures of selection across genes associated with vascular development, life-growth, adaptation to salinity and changes in flowering time, thus finding compelling evidence for a genomic basis of island syndrome in Darwin’s giant daisy radiation. This work advances understanding of factors influencing subgenome divergence in polyploid genomes, and characterizes the quick and pronounced genomic changes in a specular and diverse radiation of an iconic island plant radiation.

## Introduction

As naturalists set sail to explore the world, the distinctiveness of insular species stood out from the remaining biota. The collections carried out in the Galápagos, Cape Verde and Malay archipelagos were key for the development of the theory of natural selection (C. Darwin 1859; B. Y. C. Darwin et al. n.d.) and biogeography (Wallace 1962). More recently, Ernst Mayr’s work, which set the scene for the modern synthesis (Mayr 1942), focused heavily on island biota (Emerson 2008). The central role of remote archipelagos in our understanding of evolution is not coincidental. Organisms colonizing these regions encounter highly distinct microenvironments that provide abundant ecological niches and thus ideal conditions for rapid adaptive radiation (Lomolino, Riddle, and Whittaker 2017). The ‘island syndrome hypothesis’ predicts the repeated and pronounced phenotypic shifts that species may undergo after colonizing islands, as a result of novel selective pressures and empty ecological space (Baeckens and Van Damme 2020). While the island syndrome hypothesis has been well established (Burns 2019; Baeckens and Van Damme 2020), its integration with genomic evidence still lags. For instance, while body size differences in animal lineages are the textbook example of an island syndrome (e.g., pygmy mammoths and giant tortoises), the extent to which these changes are hereditary (genetic) or induced by different food sources (diet) has yet to be documented for many lineages. Considering the rapid and drastic changes characteristic of these radiations, it can be expected that rearrangements in genome structure contribute to the adaptation to novel environmental conditions.

Because the most prominent examples of adaptive radiation and island syndromes feature animal lineages, such as Darwin’s finches, our understanding of these phenomena in plant lineages lags (Burns 2019). As plants colonize archipelagos, they typically and repeatedly undergo shifts in leaf morphology, dispersal ability, lifespan and size (Burns 2019). This is exemplified by the daisy family (Asteraceae) which has radiated in Hawai’i (*Bidens* radiation and silversword radiations) (Knope et al. 2012; Baldwin and Sanderson 1998; Knope et al. 2020), Macaronesia (*Sonchus, Tolpis, Argyranthemum* and *Cheirolophus* radiations) (S. C. Kim et al. 1996; Gruenstaeudl, Santos-Guerra, and Jansen 2013; Vitales et al. 2014; White et al. 2018, 2020), Juan Fernández (*Erigeron* and *Robinsonia* radiation), Ryukyu (*Ainsliaea* radiation) (Mitsui and Setoguchi 2012), Tristan da Cunha (*Commidendrum* and *Melanodendron* radiation) (Eastwood, Gibby, and Cronk 2004), Mauritius (*Psiadia* radiation) (Besse et al. 2003), and Polynesia (*Tetramolopium* radiation)(Whitkus 1998).

One iconic, yet understudied, plant radiation is the remarkable diversification of daisies in the genus *Scalesia* (Blaschke and Sanders 2009; Fernández-Mazuecos et al. 2020; Crawford et al. 2009; U. Eliasson and U 1974). This genus consists of ca. 15 species, which have colonized moist forest, littoral, arid, dry forest, volcanic soil, lava gravel and fissured environments across varied elevations (Itow 1995; Blaschke and Sanders 2009). This adaptive ability has been linked to *Scalesia*’s exceptional variation in leaf morphology, which may be associated with the adaptation to dry and humid environments (Stöcklin 2009; Fernández-Mazuecos et al. 2020), capilum morphology and habit. The outstanding phenotypic and ecological variation has led previous authors to refer to it as the ‘Darwin finches of the plant world’ (Stöcklin 2009). All *Scalesia* species are ancestrally tetraploid (2*n*=4*x*=68) (Ono 1967; Uno Eliasson 1974), and the polyploid genetics may have provided the genetic grist for adaptive radiation, as has been speculated for other island floras (Meudt et al. 2021).

Here, we describe a high-quality chromosomal reference genome assembly and annotation for *Scalesia atractyloides*. This species was chosen since it is a critically endangered species and because it belongs to the most basal lineage in the endemic radiation (Fernández-Mazuecos et al. 2020). A chromosome-resolved assembly has allowed us to identify and separate the two ancestral genomes that united in the polyploidization event, and to compare gene and transposable element distribution across and between these subgenomes. Annotation of genes using PacBio IsoSeq RNA afforded a high-quality annotation of the genome, and the detection of selection and gene-family expansions that implicate the genomic basis for island syndrome traits in plants.

## Results and discussion

### Genome assembly, annotation, and quality control

*The Scalesia atractyloides* genome assembly is of remarkably high contiguity (Figure 1A), consisting of 3,216,878,694 base pairs (3.22 Gbp) distributed over 34 chromosome models, in line with previous cytological evidence (Spring, Heil, and Vogler 1997; Uno Eliasson and Others 1974; Schilling, Panero, and Eliasson 1994). The N_90_ was of 31, corresponding to all but the three smallest chromosomes (n =34) and LN_90_ was 81.66 Mbp. Flow cytometry estimates (Supplementary Information; Supplementary Table 01), however, suggest a genome size of ca. 3.9 Gbp, and thus ∼700 Mbp were likely collapsed by the assembler or removed by *purgehaplotigs* (Peona, Weissensteiner, and Suh 2018). Despite this likely collapse of repeats, we were able to annotate 76.22% of the genome as repeats, which were masked by *RepeatMasker* (∼2.5 Gbp). Considering the whole genome, 47.9% of the genome was composed of long terminal repeat (LTR) retroelements, of which 16.2% were Copia and 31.54% Gypsy elements (Supplementary Information; Supplementary Table 2), and 26.32% were unclassified repeats.

**Figure 1.**
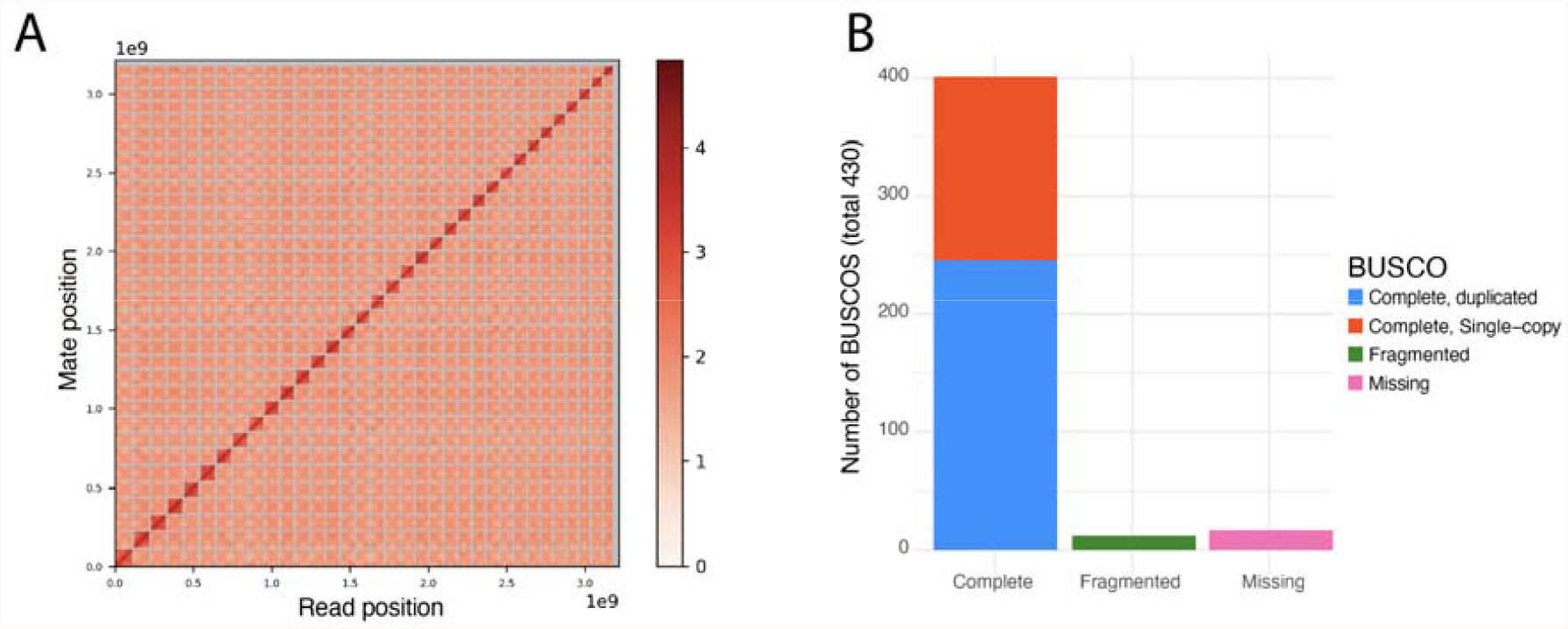
Chromosome-resolved assembly of *the Scalesia atractyloides* nuclear genome. A) Link density histogram, with 34 linkage groups (chromosome models) identified by contiguity ligation sequencing. The x and y axes show mapping positions of the first and second read in read pairs. B) Viridiplantae BUSCO set, which offers a characterization of universally conserved orthologue genes.

The IsoSeq transcriptome recovered 46,375 genes and 224,234 isoforms (Supplementary Information; Supplementary Figure 01). Using this as evidence and *ab initio* models, we retrieved 43,093 genes from the annotation. Of the 430 Viridiplantae odb 10 BUSCO groups used in a search of the genome (Figure 1B), 401 were found as complete (93.3%), of which 245 were found as duplicate (57%), 156 as complete and single-copy (36.3%), and 12 as fragmented (2.8%). Only 17 were absent (3.9%). When running OrthoFinder including Scalesia and five other Asteraceae chromosome-resolved assemblies, we found that 34% of all the orthogroups included genes from the five genomes. This overlap indicates a high-quality gene annotation (Supplementary Information; Supplementary Figure 02). The proportion of annotated repeats and number of genes is within the variation reported for Asteraceae. For instance, the assembly of the closely related sunflower (*Helianthus annuus*) reference genome includes ∼52,000 protein coding genes and has a repeat content of 74% (Badouin et al. 2017).

### Subgenome identification & evolution

The identification of subgenomes was carried in two steps. On the first step, we assigned the 34 chromosomes into 17 homeolog pairs by identifying and mapping duplicated conserved orthologous sequences (COS; Supplementary Information; Supplementary Table 03; Figure 2 A). This step only allowed the identification of pairs (homeologs), and did not allow the assignment of subgenome identity within pairs. Homeolog exchanges were therefore not a concern here (Edger et al. 2018). On the second step, we used the *k*-mer spectrum to identify ‘fossil transposable elements’, which are transposable elements that were replicating while both subgenomes were separate (i.e. after the speciation event, before the polyploidization event). Since different genomes accumulate different transposable elements, we hypothesized that some transposable element families in either subgenome will maintain frequency biases (Session et al. 2016; Mitros et al. 2020). In short, transposable element families active before the divergence of the two parental lineages are predicted to be approximately equally represented in either subgenome, whereas elements activated after the divergence of the parental species are predicted to be differently represented on the Scalesia subgenomes. Using the *k-mer* spectrum, we selected k-mers that were in high numbers (i.e. repeats/TEs) and unevenly represented between chromosome-pairs identified in the previous step (i.e. active during the separation period). Using this selection of *k-mers* we ran a hierarchical clustering approach that grouped chromosomes into two groups (two subgenomes; Supplementary Information; Supplementary Figure 03). To confirm this assignment, we explored the output from *RepeatMasker*, finding transposable element families unevenly represented across subgenomes (Figure 2B), as predicted. The identification of differently represented transposable element families also provides compelling evidence that the *Scalesia* radiation is of allopolyploid lineage. Island floras are characterized by a high degree of paleo-allopolyploids, where the variation brought forth by ploidy may underpin the diversification to multiple environments (Julca et al. 2020; te Beest et al. 2012) -a scenario which is line with the evolutionary history of *Scalesia*.

**Figure 2.**
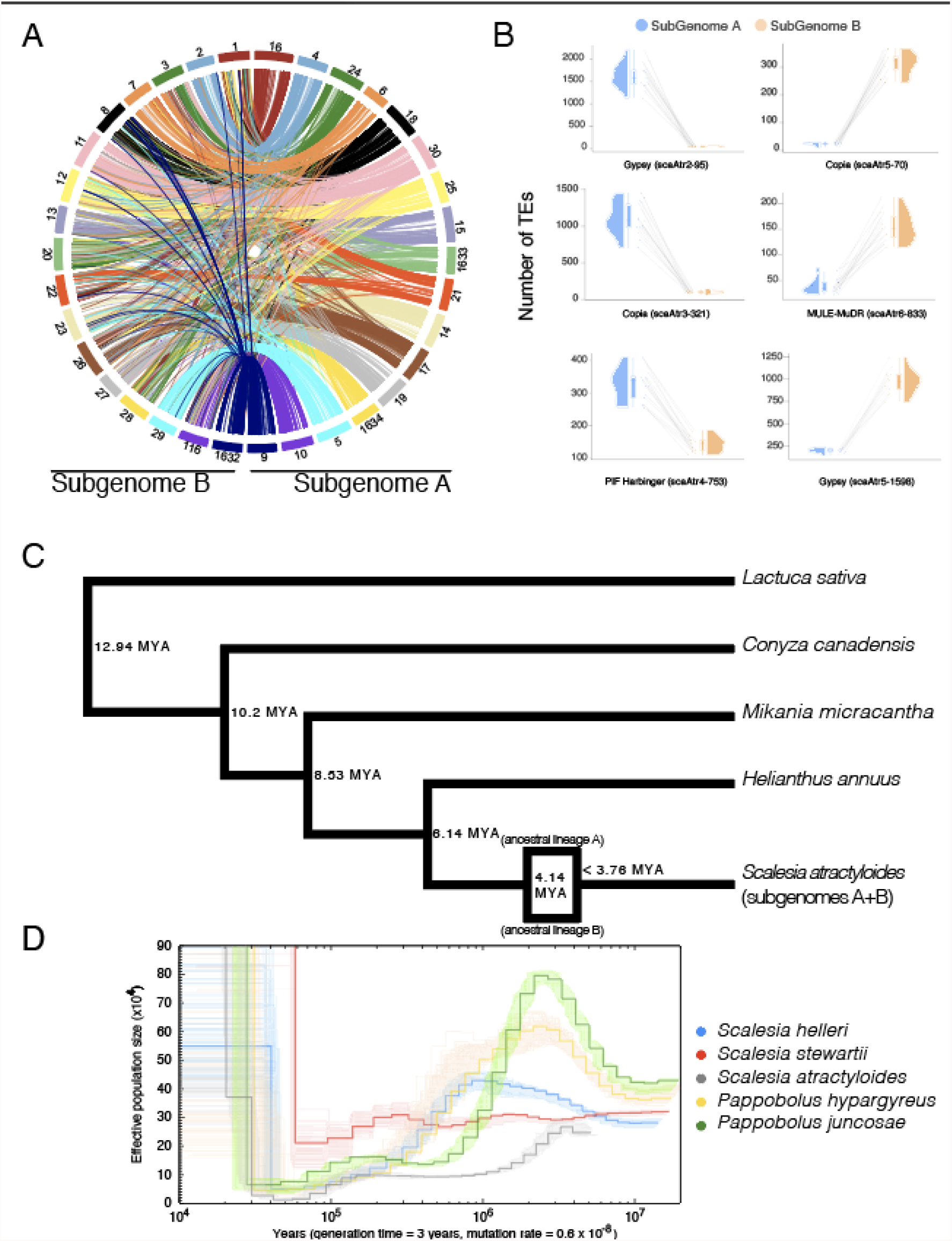
Subgenomes and evolutionary history of *Scalesia*. A) Circos plots displaying the 34 pseudo-chromosomes in the assembly. Pairs are organized to the left and right from the top, and have the same colour-coding; B) Families of transposable elements (TEs) that are differently represented on each subgenome. These TE families were likely active while the two subgenomes were separated and thus confirm subgenome identification. Each data point corresponds to a chromosome in a subgenome (subgenome A in blue and B in orange). Chromosome pairs are linked by grey lines; C) Single-copy ortholog phylogeny of the studied Asteraceae genome assemblies. Node ages are provided to the right of each node, as well as the predicted time for the polyploidization event. D) Pairwise sequentially Markovian coalescent (PSMC) estimation of the demographic history of *Scalesia atractyloides*, two other *Scalesia* species, and two members of the *Pappobolus* genus, which is the sister taxon to *Scalesia*.

Using four other chromosome-level assemblies from Asteraceae (*Helianthus annuus, Conyza canadensis, Mikania micrantha, and Lactuca sativa*) and the two subgenomes, we estimated groups of orthologous genes using OrthoFinder. We obtained 710 orthogroups in which each genome had only a single member, tolerating no missing data, and used this data to construct a phylogenetic tree. The tree topology agrees with the placement of the Asteraceae lineages from a recent and comprehensive set of genomic analyses (Mandel et al. 2019). Dating of this tree was done by constraining the node separating the *Scalesia* subgenomes and *Helianthus* as 6.14 Mya (Figure 2C), following recent literature (Mandel et al. 2019). This suggests that the subgenomes diverged from their MRCA roughly 4.14 Mya, but the separation of the ancestral lineages only lasted ∼0.5 My, as calculated by LTR-family divergence (Supplementary Information; Supplementary Figure 04). Specifically, the ancestral genomes reunited in a single polyploid genome at least 3.76 Mya (Figure 2C; Supplementary Information; Supplementary Figure 04). These dates are concordant with the *PSMC* analysis which roughly indicate that the three *Scalesia* species had concordant population sizes of 250,000-300,000 circa 4 MYA (Figure 2D). Mismatches between the three genomes could result from variation in generation time in *Scalesia* (see results and discussion below), and bottlenecks suffered by populations as a result of climatic shifts in the Galapagos (Whittaker, School of Geography Robert J Whittaker, and Fernandez-Palacios 2007). These estimates are concordant with a recent dating analysis that estimated the divergence between *Pappobolus* and *Scalesia* occurred ∼3 Mya (Fernández-Mazuecos et al. 2020).

The identification of subgenomes allowed comparing the genes and transposable element distribution across chromosome pairs. We find that gene density is highest near the telomeres on both subgenomes, while transposable elements are more evenly distributed throughout chromosomes (Figure 3A). This even distribution of transposable elements is different from most other vascular plants, in which transposable element load is highest near the center and decreases towards the ends of the chromosome, and rather is reminiscent of observations in bryophyte genomes (Diop et al. 2020; F.-W. Li et al. 2020; Lang et al. 2018). Even distributions of transposable elements were also observed in the sunflower genome (Badouin et al. 2017), which may be indicative of particular transposable element regulation in the Heliantheae.

**Figure 3.**
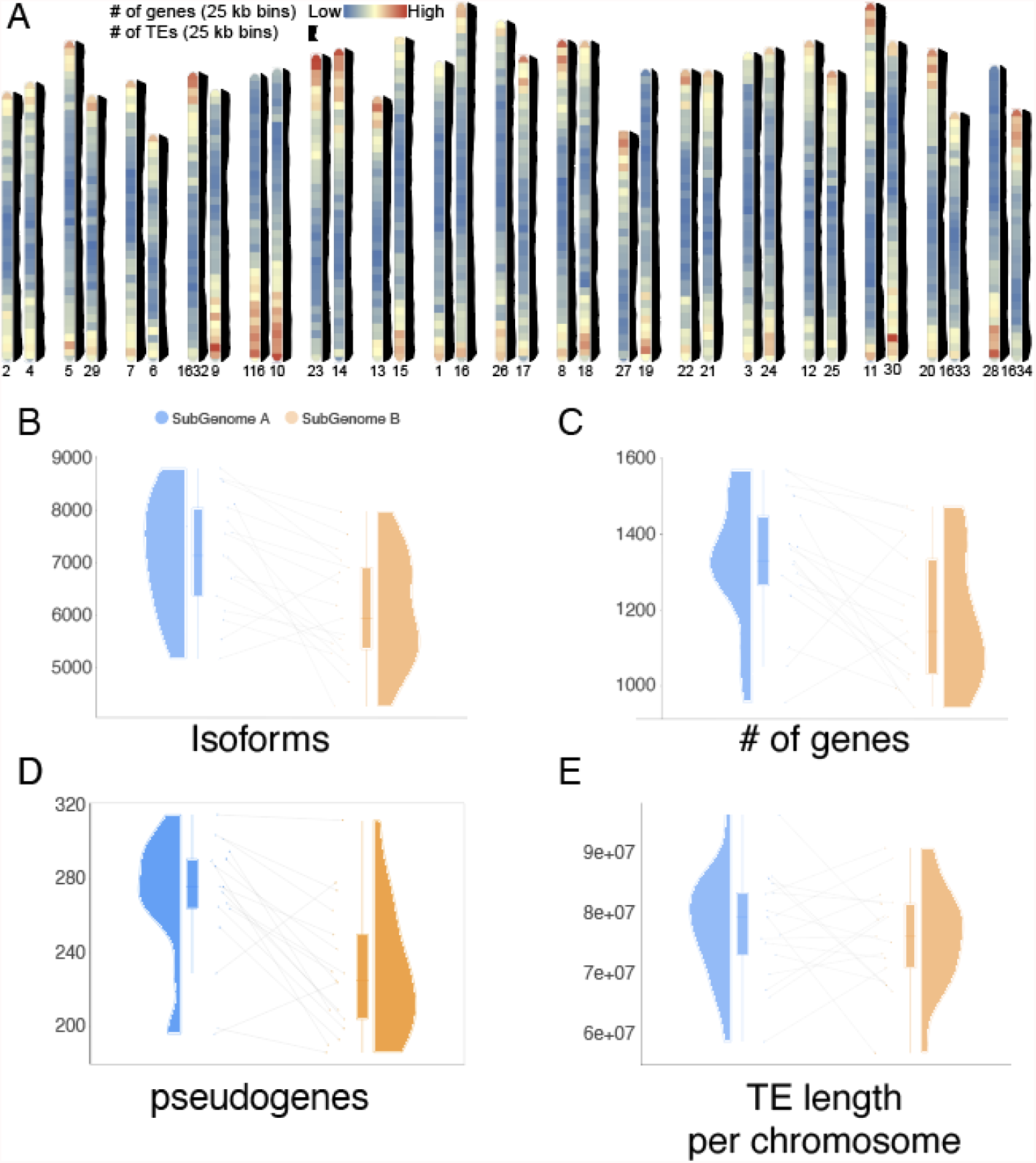
Subgenome evolution and characterization. A) Ideogram with gene and transposable element distribution in 25-kbp bins. Gene density is plotted in chromosome representations and transposable element distribution is plotted to the side of each chromosome in black. Chromosomes are arranged in homoelogous pairs. B) Number of isoforms detected for each subgenome. Each data point corresponds to a chromosome in a subgenome (subgenome A in blue and B in orange). Chromosome pairs are linked by grey lines; C) Number of genes detected for each subgenome; D) Number of pseudogenes detected for each subgenome; E) Length of transposable elements detected for each subgenome.

As two genomes unite to form a single hybrid genome, an accommodation of the two subgenomes, the process of ‘diploidization’ takes place (Bird et al. 2018; Freeling, Scanlon, and Fowler 2015; Wolfe 2001). This process can occur very quickly, with changes in transcription between subgenomes observed in 2-3 generations (Bird et al. 2021), and result in pronounced changes in gene numbers. Whereas subgenome dominance in gene expression and retention has been documented paleopolyploid plant genomes (Alger and Edger 2020; Renny-Byfield et al. 2015; Douglas et al. 2015), *Scalesia* subgenomes contain roughly equal gene and isoform contents (Figure 3 B, C), as well as pseudogene numbers and transposable element load (Figure 3 D, E). In addition to this, when running the Viridiplantae BUSCO set for each subgenome separately, we find 82.7% complete BUSCOs on subgenome A (76.6% single-copy, 6% duplicates), and 81.9% complete BUSCOs (77% single-copy, 4.9% duplicates) on subgenome B. Finally, both subgenomes are roughly the same length (subgenome A = 1,629,251,263 bp; subgenome B = 1,554,170,668 bp), and have retained the same number of chromosomes (Figure 3A). This indicates that during the past ∼3.76 million years, during which the two subgenomes have been unified in the same organism, there has not been a drastic rearrangement of either subgenome, despite a smaller accumulation of genes and pseudogenes on subgenome A. To explain this, we speculate that *Scalesia*’sadaptation to insular environments has benefitted from the genetic variation and diversity stemming from the allopolyploidization event.

### Fast evolutionary rates in Heliantheae

To further dissect the mode and tempo of polyploid subgenome evolution, we used *Synolog* (Catchen, Conery, and Postlethwait 2009) to create chromosome stability plots that would allow us to detect translocations and inversions and to quantify their impact (Figure 4). This method establishes clusters of conserved synteny by identifying single-copy orthologs shared between two genomes via reciprocal BLAST searching between all annotated protein-coding genes. From the identified synteny clusters, we calculated statistics on the orientation (forward/inverted) and chromosome location. We thereby classified genes into four categories: “Forward pair” (FP; i.e. not inverted, and the single-copy orthologs are in chromosomes from the same pair), “Inverted pair” (IP; i.e. inverted, and the single-copy orthologs are in chromosomes from the same pair), “Forward translocated” (FT; i.e. not inverted, and the orthologues are not in chromosomes from the same pair), “Inverted translocated” (IT; i.e. inverted, and the orthologues are not in chromosomes from the same pair). Comparing the two *Scalesia* subgenomes, we found 2,284 FP genes (17%), 1,760 IP genes (13%), 3,539 FT genes (27%), and 5,641 IT genes (43%), totalling 13,224 genes included in the analysis (Figure 4B). Thus, the majority of the genes have been translocated (70%), and/or inverted (56%). Interestingly, there is a mismatch between the length of these regions in the genome and the proportion of genes they contain. Specifically, we classified 434.6 Mbp as FP (23%), 189 Mbp as IP (36%), 693,2 Mbp as FT (31%), and 586,5 Mbp as IT (10%). These results are in line with the inference of rapid rates of chromosomal rearrangements in the Asteraceae based on comparisons of the sunflower and lettuce genomes (Badouin et al. 2017). The discordance between the fraction of genes, and the fraction of genome length (e.g., IT includes 43% of the genes but only occupies 10% of the genome; 46% of the subgenomes is found as inverted, but contains 56% of the genes) implicates inversions as having influenced the retention of duplicated genes after formation of *Scalesia*’s allopolyploid ancestor.

**Figure 4.**
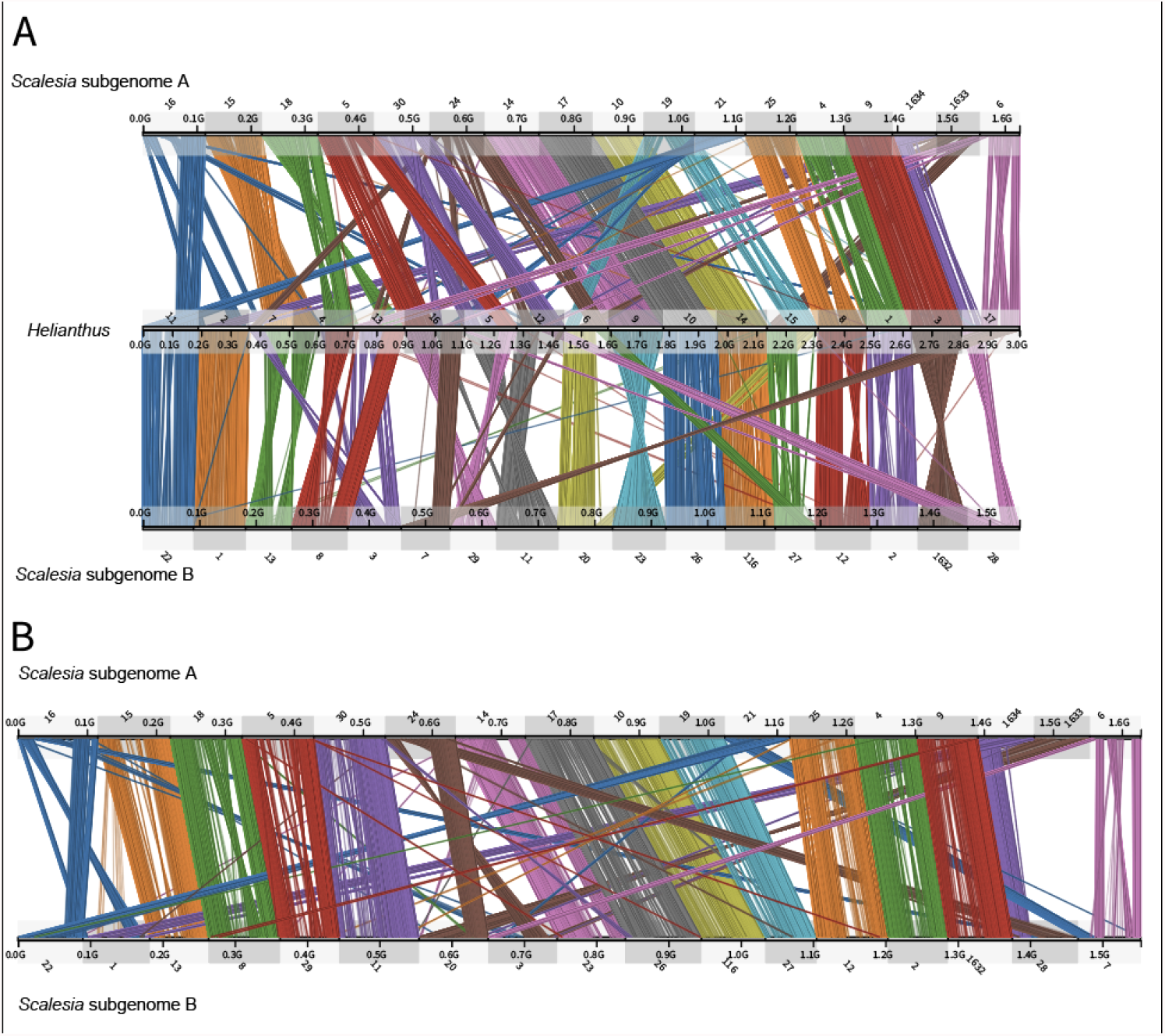
Chromosome stability plots reveal the role of inversions and translocations in the differential in each subgenome. A) Chromosome stability plot between the two Scalesia subgenomes and the *Helianthus annuus* genome. Each line connects a pair of orthologous genes, colour-coded by chromosome pair. B) Chromosome stability between the two *Scalesia* subgenomes. Each line connects orthologous genes in subgenome A and B, colour-coded by chromosome pair.

Despite its polyploid ancestry, the *Scalesia* genome has roughly the same length (3.22 Gbp) as that of the diploid sunflower (3.6 Gbp assembly). Further, *Scalesia’*s annotated gene model number (43k) is smaller than the number of gene models annotated in the sunflower genome (52k) (Badouin et al. 2017). The sunflower has 17 chromosome pairs, whereas the tetraploid *Scalesia* has 34 chromosome pairs. This indicates that *Scalesia* has likely undergone a reduction in genome length and gene numbers without a reduction in chromosome number. This evidence is consistent with hypothesized genome miniaturization in island species (J. Suda, Kyncl, and Jarolímová 2005). We cannot, however, rule out a scenario in which *Helianthus* has increased its number of genes and genome length.

### Evolutionary history of Scalesia & evidence for island syndromes

We identified 920 genes under selection (*p* < 0.05) in the *Scalesia* genome, after correcting dN/dS ratios using a Holm-Bonferroni FDR correction. To understand their function, we extracted the functional annotation using a Gene Ontology (GO) term enrichment analysis, and the results were visualised using *Revigo*. GO classifications relating to many metabolic processes under selection (Figure 5A, orange group), cellular reorganization (green group), DNA repair (yellow group), response to protein folding (maroon group), and regulation (regulation of metabolic processes, translation, gene expression, translation, nuclear division, chromosome segregation, among others; pink group; Figure 5A; Supplementary Information; Supplementary Tables 04-6). Genes inferred to have evolved under positive selection are also associated with meiosis, chromosome arrangement, and chromatin status (meiotic cytokinesis, establishment of chromosome location, chromosome separation, and chromosome segregation, among other GO classifications; Figure 5A, Supplementary Information; Supplementary Tables 04-6), and this may indicate selection at genes associated with the coexistence of two genomes.

**Figure 5.**
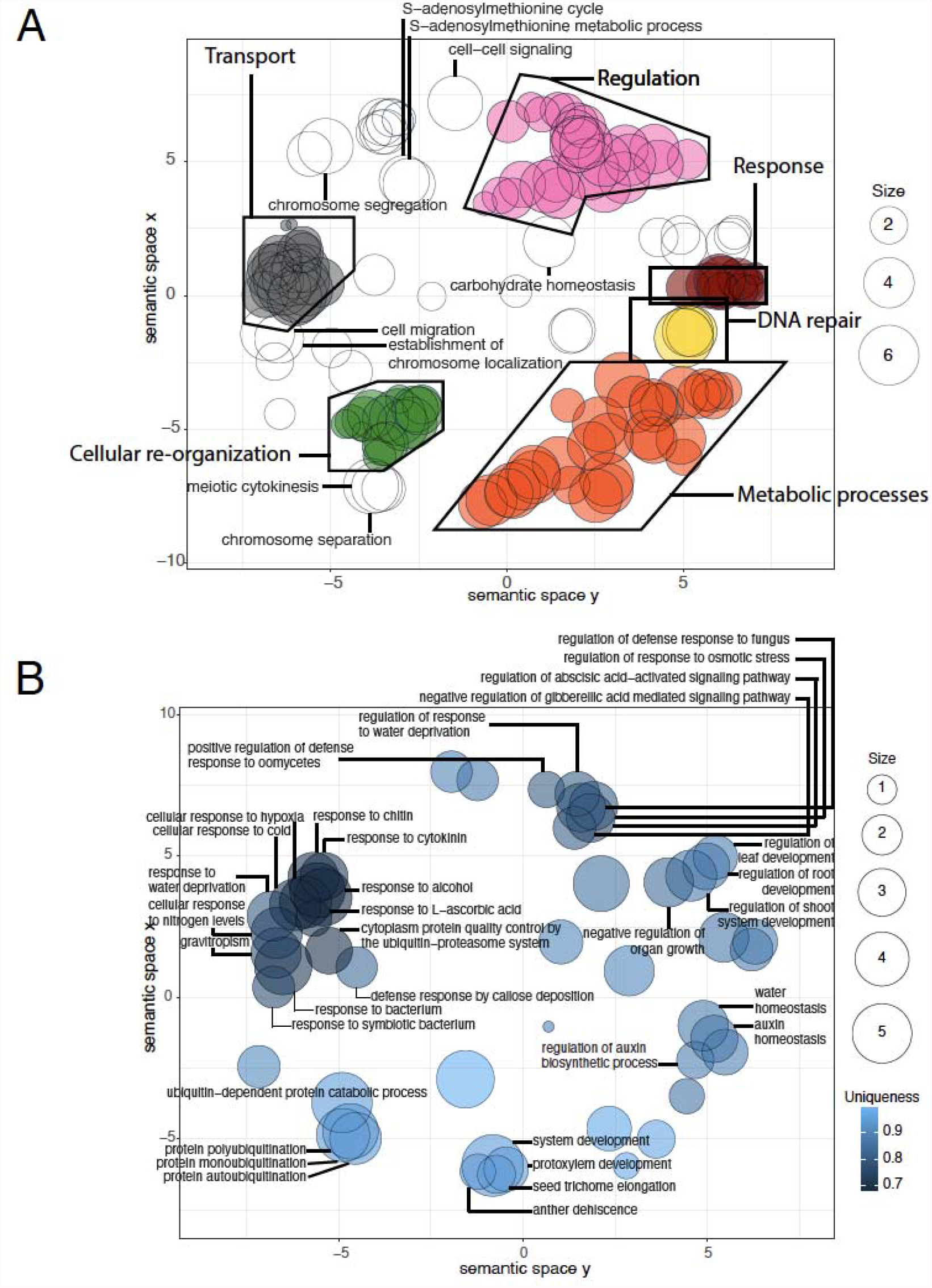
Positive selection and gene family expansion across the *Scalesia atractyloides* genome. A) GO term enrichment of the genes under selection across the genome. GO terms assigned to at least four genes are labelled. Size refers to the number of genes associated with a particular GO term. B) GO term enrichment of the genes belonging to expanded gene families across the genome according to a *CAFE* analysis. Only GOs within a group of three or more overlapping circles are included. Uniqueness measures the degree to which a particular GO term is distinct relative to the whole list.

To chart the landscape of *Scalesia*’s adaptive revolution, we randomly selected 100 genes under selection and performed a systematic investigation of homolog function in *Arabidopsis thaliana* literature. The classification of these genes into non-mutually exclusive categories showed 23 genes implicated in photosynthesis and leaf morphology/metabolism, 17 in stress response, 15 in fertility and reproductive organ function, 13 in life-phase transition and growth, 11 in the embryo, 10 in root functions, eight in immunity, seven in cellular functions (mitochondrial functions, cytoplasm, DNA repair, nucleus, cell elongation or autophagy), five in meiosis/mitosis, three in the vascular system, two in general metabolism, and one in methylation (Supplementary Information; Supplementary Table 07). As leaf morphology was an important innovation in the *Scalesia* radiation, it is particularly interesting that we found selection on potential regulators of leaf morphology, including genes well known to determine leaf cell number in *A. thaliana* (E2F1) (Őszi et al. 2020; Berckmans et al. 2011), cell fate in leaves (YABBY5) (Kojima et al. 2011; Husbands et al. 2016), leaf senescence (RANBPM, LARP1C, PEN3) (Crane et al. 2019; Fu et al. 2019; Zhang et al. 2012), leaf variegation (THF1) *(Ma et al. 2015; Z. Wang et al. 2016)*, and leaf growth (PAC) (Jörg Meurer et al. 2017; Holding 2000; J. Meurer et al. 1998). It is also interesting to note that many *Scalesia* genes under selection are affected by light stimulus. A STRING analysis showed that there was selection at multiple points in the light regulatory pathways including responses to R/FR and blue light responses. These include an inhibitor of red and far-red light photoreceptor (PHL) (Endo et al. 2014, 2013), a lysine-tRNA ligase that regulates photomorphogenic responses (G. Li et al. 2011), an amino acid aminotransferase-like PLP-dependent enzymes superfamily protein that is regulated under light conditions and is associated with the photorespiration process (Basset et al. 2004; Smeekens 2006), and genes for which knock-out mutants experience alterations in light reception (DJC69, COX15) (Oravecz et al. 2006; Dal Bosco et al. 2004). This may underlie the natural history observations where *Scalesia* individuals growing in the absence of permanent light conditions show substantially retarded growth (Lawesson 1988) or higher mortality (Rivas-Torres, *unpublished data*).

Many of the stress-response genes under selection in *Scalesia* are associated with osmotic stress in *A. thaliana*, concomitant with evidence that the *Scalesia atractyloides* habitat is characterized by arid conditions such as the Galápagos’ arid zone, the littoral zone, and fissured lava areas (Blaschke and Sanders 2009; Itow 1995). For instance, we found five genes (“Leucine-rich repeat protein kinase family protein”, MPPBETA, “leaf osmotic stress elongation factor 1-β-1”, AT2G21250, VAP27-1) associated with osmotic stress (Tan et al. 2010; J. Y. Kim et al. 2007; ten Hove et al. 2011; Jakoby et al. 2008; Fox et al. 2020; P. Wang et al. 2019), but also heat shock proteins (Valverde, Groover, and Romero 2018), and regulators of stomatal closure (THF1) (Ma et al. 2015; Z. Wang et al. 2016) under selection. Other stress-associated genes under selection include those involved in response to high irradiation (ZAT10, AT1G06690, DDB2) (Bittner, Hause, and Baier 2021; Kuki et al. 2020; Czarnocka et al. 2020; Kleine et al. 2007; Castells et al. 2011; Lahari et al. 2018).

Some genes under selection are associated with growth and transitions between life stages. *Scalesia* plants’ fast rates of growth have earned them the name ‘weedy trees’, and these genes may regulate these plants’ exceptionally fast growth and tree-like habits. We find three genes under selection that cause the transition between embryonic and vegetative traits (RING1A, SWC4, ABCI20) *(A. Kim et al. 2020)*, and four genes that regulate flowering time in *A. thaliana* (ELF8, RING1A, Short-Vegetative-Phase, NRP1) (D. Chen et al. 2010; Shen et al. 2014; J. Li et al. 2017; An et al. 2020; Gómez-Zambrano et al. 2018), and height or size of the plant (CLAVATA, GH9C2, ELF8, NSL1, TUA6) (Glass et al. 2015; Markakis et al. 2012; Noutoshi et al. 2006; Fukunaga et al. 2017; Singh et al. 2021; Fal et al. 2017; He 2004; Hoson et al. 2014; Xiong et al. 2013; Whitewoods et al. 2020).

Genes under selection include some genes associated with increased sensitivity to bacteria and fungi (JAM3, ABCG16, NMT1, AT5G05790) (Kapos et al. 2015; Kessler et al. 2010; Swain, Singh, and Nandi 2015; Sasaki-Sekimoto et al. 2013; Ji et al. 2014) as well as WRKY70,which is central to immunity in *A. thaliana (Noh et al. 2021; H. Liu et al. 2021; S. Chen et al. 2021)*. These may indicate the importance of rapid evolutionary response to the new enemies and symbionts colonizing plants encounter upon their arrival to volcanic archipelagos.

Finally, we assessed the expansion and contraction of gene families in the *Scalesia* genome, finding a total of 37 significantly contracted families and 26 significantly expanded families (Figure 5B). GO enrichment testing of the expanded families uncovered significantly enriched functions associated with vascularization (secondary cell wall biogenesis, shoot system development, negative regulation of organ growth, xylem vessel member cell differentiation, protoxylem development), likely associated with plant growth in *Scalesia* (Fernández-Mazuecos et al. 2020). We also find evidence of evolutionary responses to aridity and changes in osmotic pressure in significantly expanded families (regulation of stomatal closure, response to water deprivation, response to osmotic stress, water homeostasis), similar to the genes under selection (Figure 5B). Interestingly, we detect contraction in gene families with GO terms associated with tree habits (shoot system development, regulation of organ growth, regulation of root development, xylem vessel member cell differentiation, gravitropism), adaptation to arid environments (water deprivation, stomatal closure, regulation to osmotic stress) and cold tolerance (cellular response to cold; Supplementary Information; Supplementary Tables 08-13). While this may seem contradictory, it suggests that different families have redundant functions, and the expansion of a family may lead to redundancy in another family and consequent gene loss through pseudo-gene formation.

## Conclusions

In this study, we were able to elucidate patterns of genome evolution in the critically endangered Darwin’s giant daisy tree (*Scalesia atractyloides*) by attaining a chromosome-resolved genome and by subsequently identifying two ancient genomes underlying its polyploid state. We found that both subgenomes retain a relatively similar number of genes as well as other genetic features, such as pseudogenes and transposable elements, which lead us to speculate on the role of insular evolution underlying these changes. Moreover, we uncovered the role of inversions in gene accumulation, suggesting these have played an important role in the maintenance of genes in subgenomes, and found a relatively unique pattern of transposable element accumulation within flowering of plants. Ultimately, expanded gene families and genes under positive selection indicate the first and solid evidence for genomic island syndrome in a plant, revealing an underlying genomic basis of the outstanding phenotypic variation in *Scalesia*.

## Methods

### Plant material, flow cytometry, DNA extraction, library preparation and sequencing

Tissues used for the *de-novo* genome assembly and annotation were sampled from living *Scalesia atractyloides* plant P2000-5406/C2834 cultivated in the greenhouse of the University of Copenhagen Botanical Garden collections. This plant was originally germinated from a seed collected from Santiago Island. Fresh tissue was collected and flash-frozen in dry ice or liquid nitrogen and then stored at −80C for later use.

To assist with sequencing coverage strategy and to inform genome assembly, we obtained estimates of genome size using flow cytometry following (Galbraith et al. 1983). Briefly, 50 mg of freshly collected leaves from the sample material and from the reference standard (*Solanum lycopersicum* ‘Stupické’; 2C = 1.96 pg; (Jaroslav Dolezel, Sgorbati, and Lucretti 1992) were chopped with a razor blade in a Petri dish containing 1 ml of Woody Plant Buffer (Loureiro et al. 2007). The nuclear suspension was filtered through a 30-µm nylon filter, and nuclei were stained with 50 mg ml-1 propidium iodide (PI) (Fluka, Buchs, Switzerland). Fifty mg ml-1 of RNase (Sigma, St Louis, MO, USA) was added to the nuclear suspension to prevent staining of double-stranded RNA. After a 5 minute incubation period, samples were analysed in a Sysmex CyFlow Space flow cytometer (532 nm green solid-state laser, operating at 30 mW). At least 1,300 particles in G1 peaks were acquired using the FloMax software v2.4d (Jan Suda et al. 2007). The average coefficient of variation for the G1 peak was below 5% (mean CV value = 2.72%). The holoploid genome size in mass units (2C in pg; *sensu* (Greilhuber et al. 2005) was obtained as follows: sample 2C nuclear DNA content (pg) = (sample G1 peak mean / reference standard G1 peak mean) * genome size of the reference standard. Conversion into base-pair numbers was performed using the factor: 1 pg = 0.978 Gbp (J. Dolezel et al. 2003). Three replicates were performed on two different days, to account for instrumental artefacts.

The commercial provider Dovetail Genomics extracted and purified high-molecular-weight DNA from flash-frozen leaf tissue using the CTAB protocol, and the concentration of DNA was measured by Qubit. For long-read sequencing, they constructed a PacBio SMRTbell library (∼20kb) using the SMRTbell Template Prep Kit 1.0 (PacBio, CA, USA) following the manufacturer recommended protocol. This library was bound to polymerase using the Sequel Binding Kit 2.0 (PacBio) and loaded onto the PacBio Sequel sequencing machine using the MagBeadKit v2 (PacBio). Sequencing was performed on the PacBio Sequel SMRT cell, using Instrument Control Software v5.0.0.6235, Primary analysis software v5.0.0.6236, and SMRT Link Version 5.0.0.6792. PacBio sequencing yielded 41,322,824 reads, resulting in a total of 197-fold coverage of the nuclear genome. For contiguity ligation, they prepared two Chicago libraries as described in (Putnam et al. 2016). Briefly, for each Dovetail Omni-C library, chromatic is fixed in place with formaldehyde in the nucleus and then extracted. Fixed chromatin is digested with DNAse I, and chromatin ends are repaired and ligated to a biotinylated bridge adapter followed by proximity ligation of adapter containing ends. After proximity ligation, crosslinks are reversed and the DNA is purified. Purified DNA is then treated to remove biotin that was not internal to ligated fragments. Sequencing libraries were generated using NEBNext Ultra enzymes and Illumina-compatible adapters. Biotin-containing fragments were isolated using streptavidin beads before PCR enrichment of each library. These libraries were then sequenced on an Illumina HiSeq 2500 instrument, producing a total of 1,463,389,090 sequencing reads.

To obtain RNA transcript sequences for annotation of the genome, we extracted RNA from five tissues (root, stem, young leaf, old leaf, and floral head) of *S. atractyloides* plant P2000-5406/C2834 using a Spectrum Plant Total RNA Kit (Sigma, USA) with on-column DNA digestion following the manufacturer’s protocol. RNA extracts from all five tissues were pooled. mRNA was enriched using oligo (dT) beads, and the first strand cDNA was synthesized using the Clontech SMARTer PCR cDNA Synthesis Kit, followed by first-strand synthesis with SMARTScribeTM Reverse Transcriptase. After cDNA amplification, a portion of the product was used directly as a non-size selected SMRTbell library. In parallel, the rest of amplification was first selected using either BluePippin or SageELF, and then used to construct a size-selected SMRTbell library after size fractionation. DNA damage and ends were then repaired, followed by hairpin adaptor ligation. Finally, sequencing primers and polymerase were annealed to SMRTbell templates, and IsoSeq isoform sequencing was performed by Novogene Europe (Cambridge, UK) using a PacBio Sequel II instrument, yielding 223,051,882 HiFi reads.

### Genome assembly and annotation

An overview of the bioinformatic methods is provided in *<u>JOSES_GITHUBPAGE</u>*. We assembled the genome using *wtdbg2* (Ruan and Li 2020), specifying a genome size of 3.7 Gbp, PacBio Sequel reads, and minimum read length of 5,000. The *wtdbg2* assembly consisted of contigs with 3.62 Gbp total length. This assembly was then assessed for contamination using *Blobtools* v1.1.1 (Laetsch and Blaxter 2017a) against the NT database, detecting a removing a fraction of the scaffolds. This filtered assembly was used as input to and *purge_dups* v1.1.2, which removed duplicates based on sequence similarity and read depth (Guan et al. 2020), reducing the assembly length was reduced to 3.22 Gbp. This assembly and the Dovetail OmniC library reads were used as input data for *HiRise* by aligning the Chicago library sequences to the input assembly. After aligning the reads on the reference genome using *bwa, HiRise* produces a likelihood model for genomic distance between read pairs, and the model was used to identify misjoints, prospective joints, and make joins. After *HiRise* scaffolding, the N_50_ increased to 16, and the N_90_ to 31, corresponding to all but the three smallest chromosomes (*n* =34), while the LN_50_ was 94.2 Mbp and the LN_90_ was 81.66 Mbp. The largest scaffold was 116.23 Mbp. In total, *HiRise* scaffolding joined 1,329 scaffolds (suppl, assemblathon script). We used the Assemblathon 2 script (https://github.com/ucdavis-bioinformatics/assemblathon2-analysis) (Bradnam et al. 2013) to assess assembly quality.

To annotate genes, we first masked repeats and low complexity DNA using RepeatMasker v4.1.1 (Smit, Hubley, and Green 2013) using the ‘Asteraceae’ repeat database with Repbase database. After this first round, we ran RepeatModeler v2.0.1 (Flynn et al. 2020) on the masked genome to obtain a database of novel elements (*denovo*). This database was subsequently used as input to *RepeatMasker* for a second round of masking the genome. To find gene models, we first assembled a transcriptome using PacBio HiFi data and following the IsoSeq3 pipeline (Pacific Biosciences). Processing of the RNA data involved clipping of sequencing barcodes (lima v2.0.0), removal of poly(A) tails and artificial concatemers (Isoseq3 refine v3.4.0), clustering of isoforms (Isoseq3 cluster v3.4.0), alignment of the reads to the reference genome using (pbmm2 align v1.4.0), characterization and filtering of transcripts (*SQANTI3* v1.0.0) (Tardaguila et al. 2018). Genome annotation was carried out using the *MAKER2* pipeline v2.31.9 (Holt and Yandell 2011; Moore et al. 2008), using a combination of *ab-initio* and homology-based gene predictions (using Asteraceae protein sets). Since no training gene models were available for *Scalesia atractyloides*, we used *CEGMA* (Parra, Bradnam, and Korf 2007) to train the *ab-initio* gene prediction software *SNAP* (Korf 2004). In addition to the *ab-initio* features, we used the PacBio-based transcriptome as a training set for the gene predictor *AUGUSTUS* (Keller et al. 2011), and as direct RNA evidence to *MAKER2*. Finally, when running *MAKER2* we specified, model_org=simle, softmask=1, augustus_species=arabidopsis and specifying *snapphmm* to training of *SNAP*. To assess the quality of the gene models we started using BUSCO and the viridiplantae odb v10 set (Simão et al. 2015; Waterhouse et al. 2018; Seppey, Manni, and Zdobnov 2019).

### Demographic reconstruction using PSMC

To complement the *S. atractyloides* genome, we generated shotgun genomic data from DNA extracts of specimens of *S. helleri* B. L. Rob. and *S. stewartii* Riley, as well as the outgroup species *Pappobolus hypargyreus* and *P. juncosae*. Briefly, the *S. helleri* and *S. stewartii* specimens were extracted with a Qiagen DNeasy 96 Plant Kit. The *P. hypargyreus* and *P. juncosae* extracts were previously reported (Fernández-Mazuecos et al 2020). DNA extracts for these four specimens were sent to the commercial provider Novogene for dsDNA library preparation, and they were sequenced on the Illumina NovaSeq platform in 150-bp PE mode. For these sequence data, we used *FastQC* v0.11.8 to check for quality of raw reads (Andrews 2017), identified adapters using *AdapterRemoval* v2.3.1, and removed them using *Trimmomatic* v0.39 (Schubert, Lindgreen, and Orlando 2016; Bolger, Lohse, and Usadel 2014). These sequences were then aligned to the *S. atractyloides* genome using the *mem* algorithm of *bwa* (H. Li and Durbin 2009), and reads with a mapping quality below 30 were removed, resulting in a final high-quality sequencing depth of about ∼15x. Alignments were then processed and analysed using *PSMC* (Heng Li and Durbin 2011). Specifically, this processing involved calling variants using the *bcftools mpileup* and *call* algorithms, considering base and mapping qualities above 30 and read depths above 5 (Danecek et al. 2021), and posterior processing of the files using *fq2psmcfa*. For the *PSMC* run we specified a maximum of 25 iterations, initial theta ratio of 5, bootstrap, and a pattern of “4+25*2+4+6”. To plot files we used the util *psmc_plot*.*pl* specifying a generation time of 3 years and a mutation rate of 6e^-9^, and constrained the y-and x-axes to 50 and 20,000,000, respectively.

### Determination of subgenomes, and testing for subgenome dominance

We reasoned that homologous chromosomes would share conserved orthologue sets (COS). We used the Compositae-COS as baits (available through github.com/Smithsonian/Compositae-COS-workflow/raw/master/COS_probes_phyluce.fasta (Mandel et al. 2014), running *phyluce* to mine for COS in the genome assembly (Faircloth 2016; Faircloth et al. 2012). This pipeline, however, is designed for single-copy COS, and we modified the python script to provide duplicates. We then constructed a matrix of COS-assignation using double-copy COS (Supplementary Information; Supplementary Table 03).

Duplicated-COS provided a solid determination of chromosome pairings but did not reveal which member of the pair belongs to either subgenome (subgenome A or B *hereafter*). To distinguish this, we analysed the *k*-mer spectrum (Session et al. 2016). We hypothesized that given a period of separation between the two subgenomes, they have accumulated different repeat content and transposable elements. To do so, we ran the software *Jellyfish* (Marçais and Kingsford 2011) for each chromosome independently, thus obtaining a chromosome-by-chromosome frequency of 13-mers. To ensure we obtained only repeats, we selected 13-mers represented only >100 times in each chromosome, and kept only *k*-mers which were twice represented on a member of each pair. Using R, we computed a distance matrix and a hierarchical clustering, which neatly separated members of each pair into two groups (Supplementary Information; Supplementary Figure 03).

To confirm the quality of subgenome assignment we took two independent approaches. First, we did a circos plot using the masked regions of the genome. To do the circos plot we aligned the masked subgenomes to each other using mummer (Delcher et al. 1999; Kurtz et al. 2004), and plotted the circos using the ‘Circos, round is beautiful’ software (Krzywinski et al. 2009). Second, we studied transposable element representation in each subgenome benefiting from the transposable element identification done by *RepeatMasker*. In specific, we obtained the list of different annotated transposable elements from *RepeatMasker* (e.g. RTE-BovB, LINE-L1, LINE-L2, Helitron, PIF-Harbinger Gypsy, Copia, CRE), and separated the families within these groups. For each family, we counted the number of elements present on each subgenome, and plotted all the families using raincloud plots (Allen et al. 2019). To visualize genes and transposable elements along chromosomes we used the R package Ideogram (Hao et al. 2020). After finding out each subgenome, we ran BUSCO separately for each subgenome as a way of understanding subgenome specific gene loss (Viridiplantae odb10 as specified above).

### Evolutionary history of the Scalesia atractyloides subgenomes and comparative genomics

We searched the literature and NCBI for chromosome-level assemblies of the Asteraceae (05 / February / 2021), downloading the genomes of the sunflower (*Helianthus annuus* (Badouin et al. 2017)), the Canada fleabane (*Conyza canadensis; (Laforest et al. 2020)*), the ‘mile-a-minute’ weed (*Mikania micrantha*; (B. Liu et al. 2020)), and the lettuce (*Lactuca sativa*; (Reyes-Chin-Wo et al. 2017)). We downloaded the *Arabidopsis thaliana* genome from Arabidopsis.org.

To obtain sets of orthologous genes, we ran OrthoFinder (Emms and Kelly 2015) on predicted amino acid sequences (faa) and coding sequences (cds). Before running this software, we selected only the longest isoforms of both files, and removed sequences with stop codons. On the amino acids file we removed sequences with lengths below 30 bp using kinfin’s filter_fastas_before_clustering.py script (Laetsch and Blaxter 2017b). We ran OrthoFinder on various combinations of the genomes, including: 1) All Asteraceae, with subgenomes separated, (*S. atractyloides* subgenomeA, *S. atractyloides* subgenomeB, *C. canadensis, H. annuus, L. sativa, M. micrantha*); 2) All Asteraceae, and the *Scalesia* genome (*S. atractyloides* (complete), *C. canadensis, H. annuus, L. sativa, M. micrantha*); 3) *A. thaliana* and subgenomes (*S. atractyloides* subgenomeA, *S. atractyloides* subgenomeB, *A. thaliana*). A representation of run 2 and its processed results using an upset plot (Conway, Lex, and Gehlenborg 2017).

To obtain a tree of the two subgenomes we ran OrthoFinder with the two subgenomes and obtained the tree of the single-copy orthologs. This tree was made ultrametric using r8s (Sanderson 2003). To date the tree, we converted branch lengths to time estimates using a calibration point of 6.14 Mya between *H. annuus* and *S. atractyloides* following recent literature (Mandel et al. 2019). To date the divergence of the subgenomes we followed the approach of (Session et al. 2016; Lovell et al. 2021; Mitros et al. 2020). Briefly, this approach has a simple assumption: before the speciation event (which separates the ancestral lineages) and after the polyploidization event (which brings the ancestral genomes together), the accumulation of transposable elements will be similar on both subgenomes. Transposable element families which are equally represented on both subgenomes will therefore represent the pre-speciation and post-allopolyploidization period. We focused on long-terminal repeats (LTRs) given their prevalence along the genome. We used LTRharvest to identify LTR elements (Ellinghaus, Kurtz, and Willhoeft 2008), and LTRdisgest to process these (i.e. annotating features such as genes inside LTRs). To find these features we downloaded various PFAM domains provided in (Steinbiss et al. 2009), and downloaded “Gypsy” and “Copia” domains from the PFAM online database. We converted the domains to HMMs using hmmconvert (Eddy 1992), and added HMMs from the Gypsy Database (Llorens et al. 2011). The identification and annotation of LTRs was done for the *S. atractyloides* and *H. annuus* genomes. Instead of using the whole LTR-element (i.e. whole transposable element including repeated regions and genes insides) we used only the LTR-region (long terminal repeat), and ran OrthoFinder to group closely related LTRs. We processed the orthofinder data selecting orthogroups which were in equal representation on both subgenomes, and that were also present in *Helianthus*. After this, we aligned the selected orthogroups using mafft, and cleaned poorly aligned regions using Gblocks (Castresana 2000; Talavera and Castresana 2007), with not stringent options (i.e. “allow smaller final blocks”, “allow gap positions within the final blocks”, and “allow less strict flanking regions”). After this, we removed sequences with more than 50% missing data, and re-checked whether numbers of TEs were still balanced between subgenomes. We then re-aligned the data using mafft and inferred a tree for each ortholog. We kept only orthogroups where the *S. atractyloides* sequences were monophyletic, and where both subgenomes were non-monophyletic. For the final set of orthogroups passing all this filtering, we calculated pairwise Jukes Cantor distance between each *S. atractyloides* LTR-region; and between each *S. atractyloides* and *H. annuus*. The Jukes Cantor distances were plotted in R and we analysed the overall frequency and converted it to million of years distance by a simple three rule with the *Helianthus* divergence with *Scalesia* of 6.14 Mya (Supplementary Information; Supplementary Figure 04).

### Signatures of selection and expanded gene regions

Using the *Scalesia* genome together with the remaining Asteraceae genomes we ran CAFE analyses (De Bie et al. 2006; Mendes et al. 2020) to estimate significant gene family expansions and contractions. Briefly, we did an all-by-all BLAST to identify orthologues in the dataset and estimated significantly expanded and contracted families using CAFÉ. To interpret the data we relied on Gene Ontology Annotation. We obtained GOs for the annotated *Scalesia* genes by means of two complementary approaches: 1) by using the Interproscan command-line version (Jones et al. 2014), using the NCBI’s Conserved Domains Database (CDD), Prediction of Coiled Coil Regions in Proteins (COILS), Protein Information Resource (PIRSF), PRINTS, PFAM, ProDom, ProSitePatterns and ProSiteProfiles, the Structure–Function Linkage Database (SFLD), Simple Modular Architecture Research Tool (SMART), SUPERFAMILY, and TIGRFAMs databases; 2) by extracting the curated Swiss Prot database from UniProt (Viridiplantae). We blasted the *Scalesia* genes to this database and kept hits with an e-value below 1e-10. We then extracted the GOs from each gene from the database and assigned these to *Scalesia*’s correspondent orthologs (e-value below 1e-10). Genes belonging to significantly expanded gene families in the *S. atractyloides* genome were analysed using a GO enrichment analysis. To do so, we used the TopGO package using the ‘elim’ algorithm which takes GO hierarchy into account (Alexa, Rahnenfuhrer, and Others 2010; Alexa and Rahnenführer 2009), this were then plotted with *REVIGO (Supek et al. 2011)*.

To test which genes are under positive selection in *S. atractyloides* genome, we retrieved the orthogroups from all Asteraceae, and aligned the cds from each orthogroup using prank (Löytynoja 2014). Considering the divergence in the genomes, as well as evidence for fast evolution in Asteraceae genomes (including this paper), we ran zorro (Wu, Chatterji, and Eisen 2012), to assess alignments. Zorro scores each alignment position with a score between 0-10, and we selected only alignments with an average score position of 5 or greater. For each of these, we inferred a tree using iqtree and ran HyPhy using its aBSREL selection test (Smith et al. 2015; Pond, Frost, and Muse 2005). To summarize these results we: 1) ran a GO enrichment (as specified above) and plotted results using *REVIGO*; 2) identified the *Arabidopsis* ortholog to each of the *Scalesia* gene under selection using BLAST, and analysed the *Arabidopsis* literature for that particular gene (Supplementary Information; Supplementary Table 07); 3) we ran a STRING analysis using the *Arabidopsis* ortholog (Szklarczyk et al. 2015), exploring the potential protein-protein interactions among genes under selection. Interaction scores of edges were calculated based on the parameters Experiments, Co-expression, Neighborhood, Gene fusion and Co-occurrence. Edges with interaction score higher than 0.400 were kept in the network. After excluding genes with no physical connection, the STRING network had 627 nodes with 470 edges (PPI enrichment p-value <0.001). To simplify the densely connected network into potential biologically functional clusters, we used the distance matrix obtained from the STRING global scores as the input to perform a k-Means clustering analysis (number of clusters = 6). 4 out of the 6 clusters are enriched for biological processes related GO terms. Cluster 1 (red bubbles) were enriched for the GO term metabolic processes, cluster 3 (lime green bubbles) for histone modifications and chromosome organization, cluster 4 (green bubbles) for response to light, and cluster 6 (purple bubbles) for ribosomal large subunit biogenesis and RNA-processing.

## Data Availability

We are in the process of submitting raw reads to ENA and the reference genome to Dryad.

## Author contributions

J.Ce. designed the experiment, processed and analysed the data and wrote the manuscript. B.P., J.M.L.G., A. R.-C., J.Ca., S.B., J.V., S.L., D.M. helped analysed the data. J.L. was responsible for flow cytometry analyses. C.K., L. S.-B. helped retrieving DNA/RNA, N. W., M.N., P.J.D.,G.R.-T. obtained permits. M.F.-M., P.V., R.M. obtained the outgroups. G.P., A.S., N.S., N.R.N., O.S., M.T.P.G., J.H. L.-M., L.R. contributed with senior expertise in data generation and interpretation, Alexander Suh22,23. M.D.M. obtained funding, supervised J.C. and wrote the manuscript. All the authors revised and approved the manuscript.

## Acknowledgements

JC is grateful to Simen R. Sandve for fruitful discussion, Martin LaForest for sharing genome annotations for his organism, and Henning Adsersen for botanical expertise and logistical support in utilizing the University of Copenhagen botanical collections. Jennifer Mandel kindly shared the Asteraceae COS. The collection and photography of specimens, and the preparation of this manuscript benefited enormously from the cooperative assistance of the personnel of the Charles Darwin Foundation Research Station, who made arrangements for collecting trips, arranged laboratory space, and offered encouragement and support throughout the project. *Scalesia* specimens were collected under the Galápagos National Park research permit number PC-001/98 PNG and MAAE-DBI-CM-2021-0213. This publication is contribution number 2426 of the Charles Darwin Foundation for the Galápagos Islands. This work was supported by the Norwegian Research Council via project number 287327 awarded to MDM.

